# Machine learning to predict the source of campylobacteriosis using whole genome data

**DOI:** 10.1101/2021.02.23.432443

**Authors:** Nicolas Arning, Samuel K. Sheppard, David A. Clifton, Daniel J. Wilson

## Abstract

Campylobacteriosis is among the world’s most common foodborne illnesses, caused predominantly by the bacterium *Campylobacter jejuni*. Effective interventions require determination of the infection source which is challenging as transmission occurs via multiple sources such as contaminated meat, poultry, and drinking water. Strain variation has allowed source tracking based upon allelic variation in multi-locus sequence typing (MLST) genes allowing isolates from infected individuals to be attributed to specific animal or environmental reservoirs. However, the accuracy of probabilistic attribution models has been limited by the ability to differentiate isolates based upon just 7 MLST genes. Here, we broaden the input data spectrum to include core genome MLST (cgMLST) and whole genome sequences (WGS), and implement multiple machine learning algorithms, allowing more accurate source attribution. We increase attribution accuracy from 64% using the standard iSource population genetic approach to 71% for MLST, 85% for cgMLST and 78% for kmerized WGS data using machine learning. To gain insight beyond the source model prediction, we use Bayesian inference to analyse the relative affinity of *C. jejuni* strains to infect humans and identified potential differences, in source-human transmission ability among clonally related isolates in the most common disease causing lineage (ST-21 clonal complex). Providing generalizable computationally efficient methods, based upon machine learning and population genetics, we provide a scalable approach to global disease surveillance that can continuously incorporate novel samples for source attribution and identify fine-scale variation in transmission potential.

**Author summary:** *C. jejuni* are the most common cause of food-borne bacterial gastroenteritis but the relative contribution of different sources are incompletely understood. We traced the origin of human *C. jejuni* infections using machine learning algorithms that compare the DNA sequences of bacteria sampled from infected people, contaminated chickens, cattle, sheep, wild birds and the environment. This approach achieved improvement in accuracy of source attribution by 33% over existing methods that use only a subset of genes within the genome and provided evidence for the relative contribution of different infection sources. Sometimes even very similar bacteria showed differences, demonstrating the value of basing analyses on the entire genome when developing this algorithm that can be used for understanding the global epidemiology and other important bacterial infections.

## Introduction

*Campylobacter jejuni* and *Campylobacter coli* are among the most common causes of gastroenteritis globally and are responsible for approximately nine million annual cases in the European Union (1,2). These zoonotic bacteria are a common commensal constituent of the gut microbiota of bird and animal species (3,4) but cause serious infections in humans. Symptoms include nausea, fever, abdominal pain, and severe diarrhoea, with potential for the development of debilitating, and sometimes fatal, sequelae (5,6). Various infection sources have been identified including animal faeces, contaminated drinking water and especially raw or under-cooked poultry and other meats (7). However, effectively combating disease requires a detailed understanding of the relative contribution of different sources to human infection.

As in many other bacterial species, *Campylobacter* populations represent diverse assemblages of strains (3,8–10). Within this structured population, some lineages are more commonly observed in particular host species (3,4,11). Because of this host association, DNA sequence comparisons of bacteria from human gastroenteritis and potential reservoir populations have potential to reveal the infection source. This has identified contaminated poultry as a major source of human infection (12,13). Based on the body of evidence including DNA sequence analysis (14), targeted interventions have been implemented, including improved biosecurity measures on poultry farms, which have halved recorded campylobacteriosis cases in New Zealand (15,16).

Extending the principal of linking source-sink populations using genotype data, methods have been developed to attribute *C. jejuni* to the likely source based on bacterial gene frequencies in potential reservoir populations (17,18). Among the most common genotyping approaches for *C. jejuni* has been multi-locus sequence typing (MLST) that catalogues DNA sequence variation across seven housekeeping genes that are common to all strains (19,20). Isolates with identical alleles at all loci are assigned to the same sequence type (ST) and those with identical sequences at most or all loci are grouped within the same clonal complex (CC). Using these data, and allele frequencies, it has been possible to probabilistically assign clinical isolates (STs and CCs) to host source using source attribution models such as the asymmetric island model implemented in *iSource* (17) and the Bayesian population assignment model STRUCTURE (18,21). Both methods have been instructive in estimating the relative contribution of a range of domestic and wild animal hosts to human infection, with poultry often identified as the principal source of human campylobacteriosis across different regions and countries (17,18,22–25).

There are two main limitations when using genotype data to for bacterial source attribution. The first is that the ability to attribute is only as good as the degree of genotype segregation. For example, in *C. jejuni* there are host restricted genotypes (3,26) that can be readily attributed to a given host source when observed in human infections, as well as ecological generalists (27,28) that have relatively recently transitioned between hosts and cannot therefore be attributed with confidence (29). While host switching potentially imposes a biological constraint on quantitative attribution models, the second limitation is far more tractable. Specifically, most current source attribution methods are subject to limitations imposed by the underlying data. Reflecting the technology of the time, MLST-based source attribution is based only on a small fraction of the genome (approximately 0.2% for *C. jejuni* (25)) and there is considerable potential for better strain differentiation using current techniques.

The increasing availability of large whole genome sequence (WGS) datasets has greatly enhanced analyses of bacterial population structure and diversity (30). However, exploiting the full information can be challenging due to variable gene content and the complexity of interpreting the short reads produced by next generation sequencing. Notwithstanding this, some studies have attempted to overcome the limited discriminatory power of MLST in attribution studies by screening WGS data to identify elements (SNPs and genes) that segregate by host (31–33). Using these host segregating markers as input data has improved the resolution of existing attribution models, including STRUCTURE, and provided information about potential infection reservoirs and the UK and France. However, using bespoke marker selection approaches with software designed for MLST data does not maximize the potential of WGS data for source attribution.

Here, we present a machine learning approach using WGS data to predict the source of human *C. jejuni* infection. This has two principal advantages over existing techniques. First, building on WGS-based machine learning source attribution approaches applied to *Salmonella enterica* and *Escherichia coli* (34,35), we take an agnostic approach to identify which machine learning tool performs best from a broad range of available algorithms. Second, we use a WGS input capture approach using data types deposited in public databases allowing the analysis of existing MLST, core-genome MLST and WGS datasets and the reuse of data for continuous updatable monitoring in a generalizable framework. Thus, we aimed to overcome limitations of the currently available methods and use the output to investigate the infective potential of *C. jejuni* strains.

## Methods

### Dataset acquisition

A total of 5,798 *C. jejuni* and *C. coli* genomes isolated from various sources and host species were available on the public database for molecular typing and microbial genome diversity: PubMLST (https://pubmlst.org/) (S1 Table). WGS data corresponded to MLST ST and CC designations as well as core genome (cg) MLST classes. The dataset was divided into training (75%) and testing (25%) sets using phylogeny-aware sorting, wherein all members of one ST were sorted entirely into either training or testing sets (S1 Table). The ST based sorting accounts for the phylogenetic non-independence of samples (36). To allow for sufficient sample sizes per reservoir population (hereafter “class”), only the five most prevalent classes for MLST and cgMLST were used (chicken, cattle, sheep, wild bird and environment). For farm animals the classes “chicken” and “chicken offal or meat” were combined to “chicken” (likewise for sheep and cattle), whilst “environment”, “sand” and “river water” were combined into “environment”, consistent with previous studies (18,37).

### Feature engineering

The allelic profiles of MLST and cgMLST were used directly. To potentially exploit the gradient of separation encoded in the sequences underlying the MLST allelic profiles, we downloaded the underlying allele sequences and encoded the nucleotides as dummy variables and k-mers (k=21) using DSK (38). DSK was also used for encoding the WGS as k-mers. Using k=21 led to a prohibitively large input vector due to the number of unique k-mers found in all genomes (109,675,176). We reduced the number of k-mers by applying a variance threshold where k-mers which were present or absent in more than 99% of the samples were discarded, reducing the numbers of unique k-mers to 7,285,583. Furthermore, we performed feature selection by testing the dependence of the source labels on every individual k-mer using the Chi-Square statistic. To avoid data-leakage we only performed the feature selection using the training data and labels to select the 100,000 k-mers with the highest score.

### Algorithm training

All machine learning and deep learning was performed in Python (for a list of all algorithms see Figure 1). The xgboost library (39) was used for the gradient boosting classifiers with all other machine learners implemented in scikit-learn (40). The hyper-parameters for each classifier were chosen using Cartesian grid search on five-fold cross-validation of the training set. The Keras library (https://github.com/keras-team/keras) was used to construct deep learning algorithms aimed at supplying a wide range of commonly used architectures. We found this to work best, empirically, given that there is no principled means of architecture selection for such models. Specifically: (i) A recurrent neural network consisting of a layer with 64 gated recurrent units, a 50% dropout layer and Rectified Linear Unit (ReLU) activation layer; (ii) A 1-dimensional convolutional network with two convolutional layers of kernel size 3 and 5 respectively and 30 filters, both followed by 50% dropout layers and a ReLU layer; (iii) A Long short-term memory network consisting of one LSTM layer with 64 units and a 50% dropout layer; (iv) A Shallow dense network with one dense layer with 64 units followed by a 50% dropout layer and a ReLU activation layer; (v) A Deep dense network with 6 dense layers starting with 128 units and halving units with each successive layer. All individual dense layers are followed by a 50% dropout layer and a ReLU layer.

**Figure 1:**
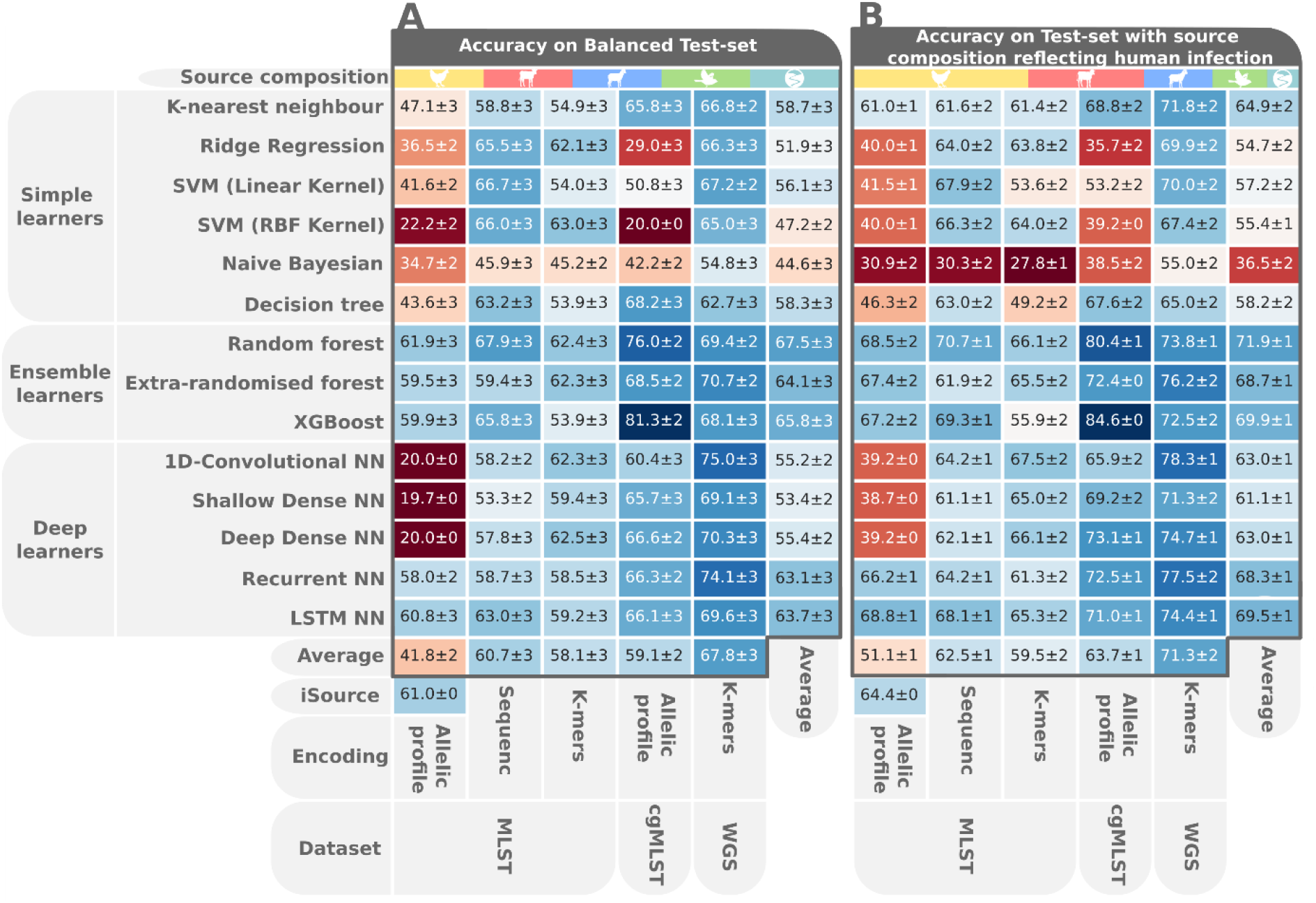
A heatmap showing classifier performance on the class balanced (A) and imbalanced (B) test set. The individual cells are coloured according to the average accuracy on 200 rounds of resampling with replacement with the variance noted next to the average accuracy. The averages of accuracy per classifiers are shown in the rightmost column, whereas the bottom column shows the averages per data type.

To all deep learning architectures, we added an output layer comprising a dense layer with soft-max activation with one unit for every class. We encoded the labels as dummy variables and used categorical cross-entropy as a loss function together with the Adam optimiser (41). Cyclical learning rates were used with a maximum learning rate of 0.1 and a minimum learning rate of 0.0001 to overcome local minima. The accuracy on the test set was measured at every epoch and the overall best performing weights were stored as a checkpoint. The data was deployed in batches of 128 samples with every batch randomly undersampled so that each class was represented in equal proportions. The training was run for 500 generations with early stopping after 50 generations.

### Algorithm testing

Both machine learning and deep learning were tested on the same 25% test set. The original data were skewed in source composition by ratios which did not necessarily reflect source origin of infection. We therefore used two methods to rebalance the classes in testing. The first test set featured an even distribution of classes, whereas the second undersampled the over-abundant chicken-origin genomes to emulate relative contribution to human disease. We used the ratios predicted by Wilson et al. (12), where *Campylobacter* genomes from chickens were 1.61 times more common than those from cattle. In both methods, rebalancing the classes was achieved by undersampling, which we repeated 200 times with replacement and averaged the accuracy over all iterations whilst also recording the variance. For performance metrics we registered accuracy, precision (positive predictive value), recall (sensitivity), F1, negative predictive value, specificity and speed. Speed was measured relative to other classifiers where a scale was defined with 0 being the slowest classifier and 1 being the quickest and all intermediate values being normalised within these confines. For comparison to previous methods, iSource was applied to the test dataset (17). Having established that XGBoost on cgMLST was the best performing source attribution method, we retrained the classifier with both training and testing data and applied it to all 15,988 human cgMLST samples available on the PubMLST database. The prediction took 892 milliseconds on a Dell OptiPlex 7060 desktop using ten threads on an Intel Core i7-8700 CPU and 16 GB RAM.

### Phylogenetic analysis

We defined the generalist index as the number of sources the ST was found in across all isolates in the dataset, which included additional samples for which only MLST data was available (S1 Table). We built a phylogeny of CC21 genomes from both source-associated and human isolates using Neighbour Joining, based on pairwise hamming distances of k-mer presence/absence in the WGS dataset, as described by Hedge and Wilson (42). We used TreeBreaker to infer the evolution of phenotypes across the phylogenetic tree of ST-21 and the most closely related sequence types. The known labels of the source-associated samples were used as phenotypic information for input into TreeBreaker (43) together with the phylogeny of CC21. TreeBreaker was run for 5,500,000 iterations with 500,000 iterations as burn-in and 1000 iterations between sampling. The phylogenetic trees were visualised with Microreact (44) and arranged alongside the results of TreeBreaker in Inkscape.

## Results and Discussion

### Machine learning outperforms popular attribution models for MLST data

In order to anchor our source attribution performance to previous efforts, we compared results using the machine learning classifiers to source probabilities estimated using the asymmetric island model implemented in iSource, which is based on MLST and the most commonly used source attribution method to date (45). The best performing machine learner on the MLST allelic profile was a random forest (61.9%/68.5% balanced/unbalanced) which performed slightly better than iSource (61%/64%) (Figure 1). Since loci within allelic profiles are deemed either to match or not, and underlying nucleotides sequences are ignored, we investigated whether exploiting the gradient of nucleotide differentiation would lead to better attribution. We used dummy variables and generated k-mers from the sequences underlying the MLST allele labels. The additional feature encodings boosted the top achieving accuracies on MLST to 67.9%/70.7% from dummy variables and 63%/67.5% from k-mers, showing the value of the additional nucleotide-level information.

### Core genome and WGS datasets increase the power of source attribution models

Having established the competitiveness of machine learning approaches for source attribution using MLST data, we turned our attention to whole genome datasets. Gene-by-gene approaches to cataloguing genomic variation in *Campylobacter* (46) and other species are a logical extension of seven-locus MLST in response to the increasing availability of large WGS datasets. Formalizing this approach to derive an approximation of the core genome for *C. jejuni* allowed the implementation of a cgMLST scheme containing 1,343 genes, that are present in the majority (>95%) of *C. jejuni* genomes (47). This has potential to increase the power of attribution models to discriminate the source of *Campylobacter* isolates based on host segregating genetic variation within the genome (37). The strong performance of tree-based ensemble classifiers continued when using cgMLST data where the XGBoost classifier achieved 81.3%/84.6% accuracy, the highest accuracy over all data types and classifiers.

q

Next, we assessed the relative performance of machine learners when applied to k-mers produced from WGS, where the average attribution performance was the highest among all datasets. The best-performing algorithm was a 1-D convolutional neural net (75.0/78.3%), performing better than the top-achieving classifier on MLST but worse than the best classifier on cgMLST despite WGS encoding more genomic information. This may be explained by the feature selection used to limit the input vector to 100,000 k-mers. Beyond comparing classifier performance on different data types, we also wanted to investigate what led to the difference in performance.

The comparison of average accuracy across all data types reveals that with an increase in encoded variation the average performance across all algorithms improves. This is especially apparent in MLST where, although capturing the same 0.3% of the genome in all isolates, the additional variation in the underlying sequences can be leveraged for better performance. When comparing the average accuracy between classifiers we observed that decision-tree based ensemble learners performed well across all datasets, with random forests performing best on average. The excellent performance of ensemble tree learners on genomic data has been reported on genomic data (48–50) and is linked to their ability to handle correlation as well as interaction of features which is an inherent feature of genomic data (50).

Amongst simple learners the K-nearest neighbour algorithm (KNN) performed best, probably owing to the hereditary nature of the phenotypic trait used as classes here. Host association is inherited both genetically, in the ability to colonise different hosts, and environmentally, in the colocation of parent and offspring cells. These patterns of inheritance result in more closely related sequences being more likely to be associated with the same phenotype. Heritability could explain the success of the KNN algorithm which is based on proximity in hyperdimensional feature space (51), which in our case is genetic similarity which is a proxy for relatedness.

The deep learners generally improved in performance with higher dimensionality of the input data - from MLST to WGS data. Among all deep learning architectures, the RNN and LSTM performed best, which was to be expected as DNA is transcribed, and mRNA translated, sequentially 5′ to 3′. Both RNNs and LSTMs process input data sequentially and input weights are also adjusted sequentially in back-propagation as opposed to the dense or convolutional architectures where input weights are tweaked concurrently. Having investigated trends across all datasets and algorithms we focused on the best-achieving classifier for a more thorough analysis of how classification performance was driven by different factors within the underlying data.

### Host transition imposes a biological limit on source attribution models

To better understand the limitations of attribution algorithms we investigated the factors driving misclassification in the different models with different datasets. The XGBoost implementation of gradient boosted decision trees, using the cgMLST dataset, was the overall best-performing classifier in our analyses. Consequently, this was used to investigate attribution performance further (Figure 2).

**Figure 2:**
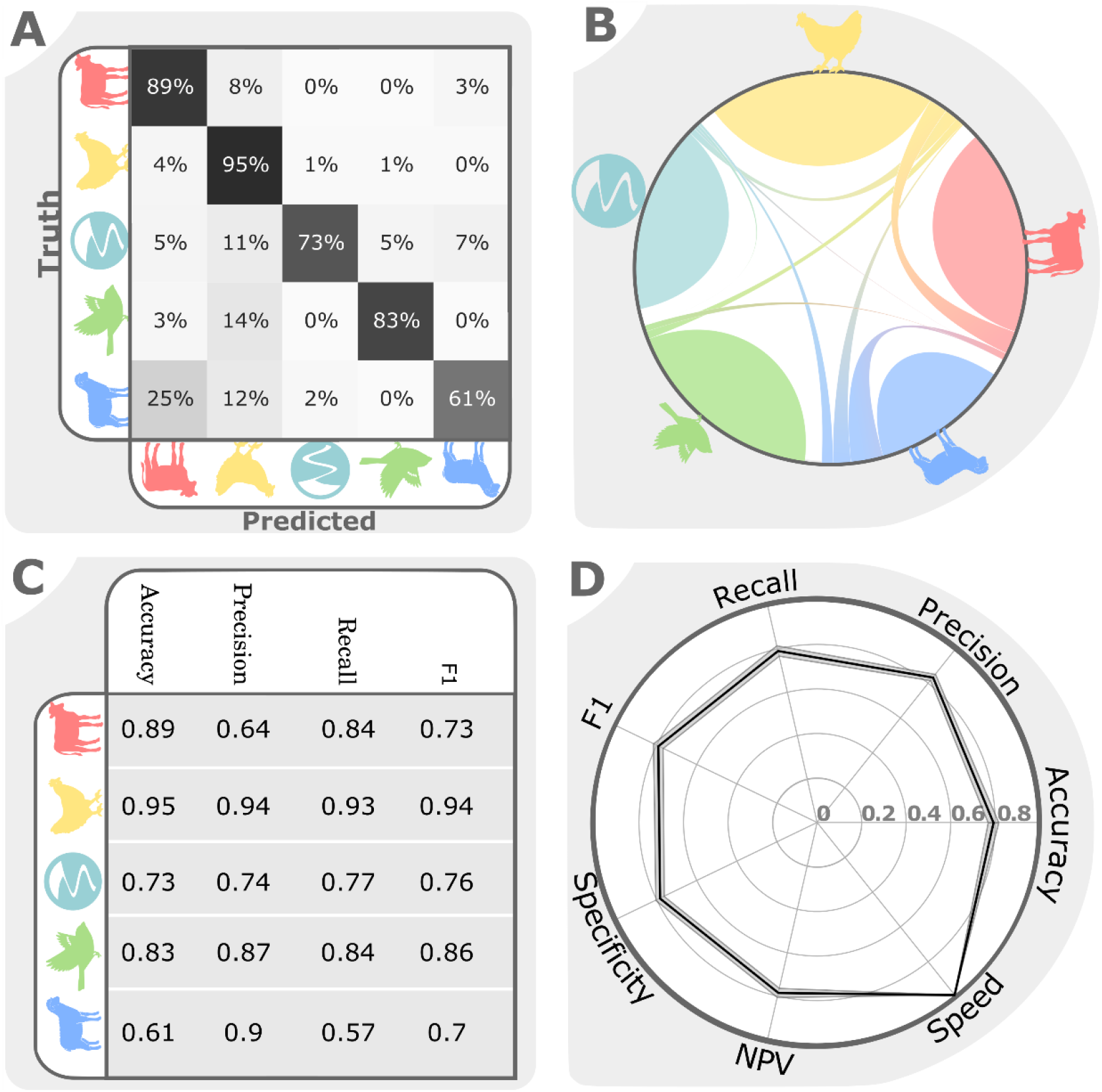
XGBoost Classifier performance on cgMLST: A) Misclassification matrix per source. The diagonal represents correct classification and off-diagonal fields are misclassifications. The percentages are calculated per row. B) Misclassification matrix as depicted in a flow diagram. C) Classifier performance on the unbalanced test set according to four different metrics per source population. D) Radar plot showing the classifier performance on the unbalanced test by seven metrics averaged over 200 rounds of resampling with replacement. The variation is depicted as a shaded surface underneath the black line representing the average.

Among all source populations the most frequent misclassification was found between sheep and cattle, which is a common source of errors in source attribution (17) owing to strongly overlapping gene pools stemming from frequent cross-species transmission that may reflect commonalities in physiological features of the ruminant gastrointestinal tracts (52). We also looked at factors besides source reservoir of the sample, as circumstances like geographical origin of the isolate (56) and the season in which they were sampled (57) have been shown to influence source attribution. We therefore stratified classification accuracy by continent, year, generalist index and *Campylobacter* species using the full non-undersampled Test dataset (Figure 3, S1 Table).

**Figure 3:**
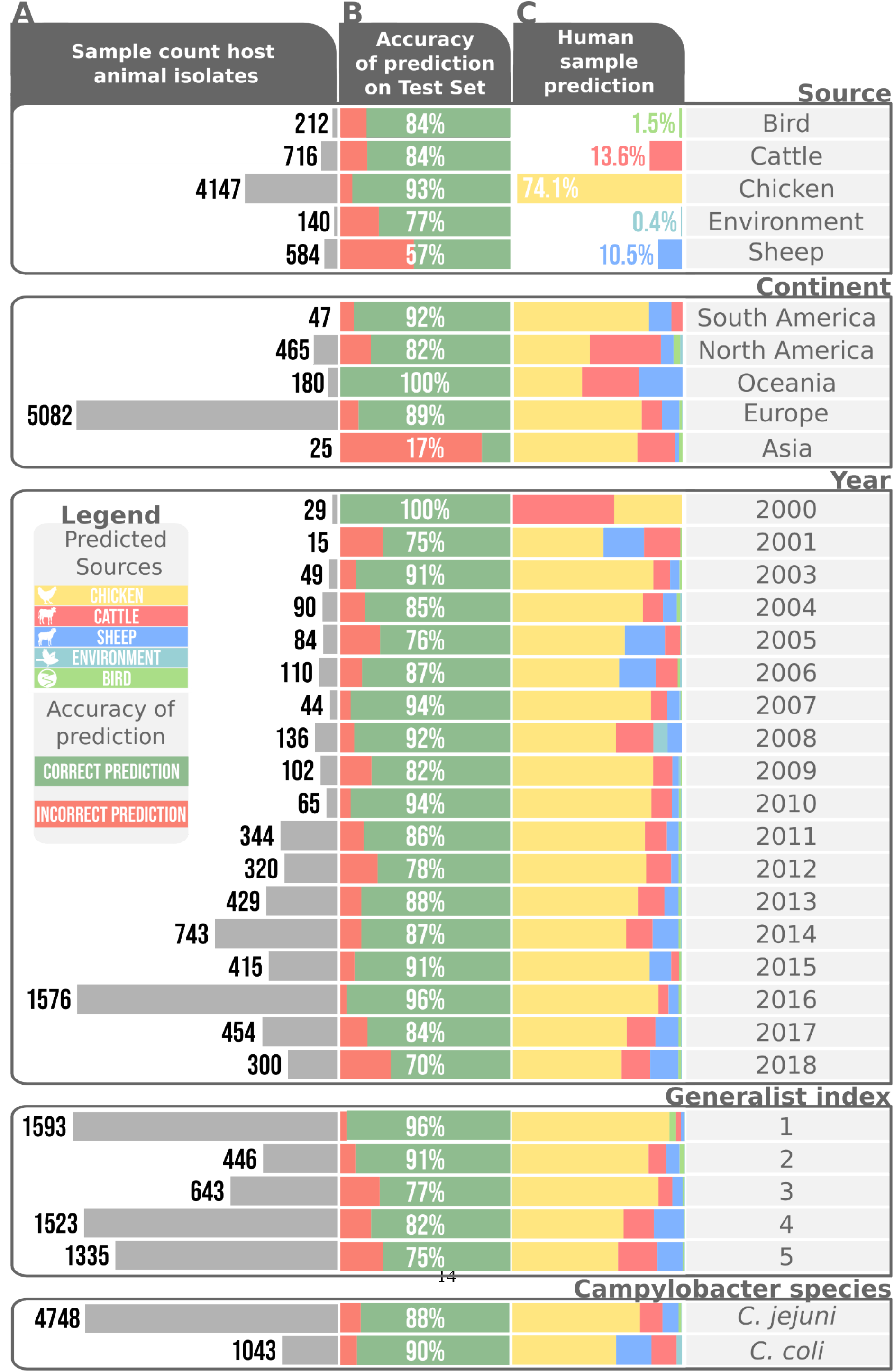
Source attribution per source, continent, year generalist index and *Campylobacter* species. A) Sample sizes across different factors in the imbalanced training set. B) Prediction accuracy on the full test dataset divided by different factors. C) Source attribution stratified into varying factors

Investigating the accuracy of the XGBoost classifier per sample size revealed that the low number of wild bird samples (212 samples; 84% accuracy) did not impede classification performance when compared to more abundant source samples like cattle (716 samples; 84% accuracy) and sheep (584 samples; 57% accuracy), presumably because wild bird STs tend to be atypical compared to the other reservoirs (46). To investigate how the ability to colonise multiple hosts affected performance, we defined a ‘generalist index’ as the number of hosts in which an ST was found across all PubMLST samples (S1 Table). The performance across generalist indices showed that strains restricted to fewer hosts were predicted with higher accuracy. This is likely due to host switching blurring the source-specific genetic signal, as previously reported (29). Consistent with this, 58% of all wild bird samples belonged to STs only found in this niche, compared to 41% in environment, 9% in cattle, 3% in sheep and 32% in chicken. Besides *C. jejuni*, an estimated 10% of campylobacteriosis cases are caused by *Campylobacter coli* (53). Consistent with previous studies, we found improved accuracy over attribution of *C. jejuni*, potentially reflecting more pronounced strain segregation by host (29), as well as a higher proportion of environmental and sheep associated strains in human infection (11,54,55) (Figure 3).

Having analysed the classification accuracy within the dataset, the machine learning method was compared to previous source attribution studies (Figure 4). Attribution of cases to chicken was consistent with higher estimates from previous studies, resulting in less attribution to all other sources, with environment identified as the source of just 0.4% of human infections. This differences in our prediction to previous studies could reflect the greater discriminatory power of cgMLST data over MLST.

**Figure 4:**
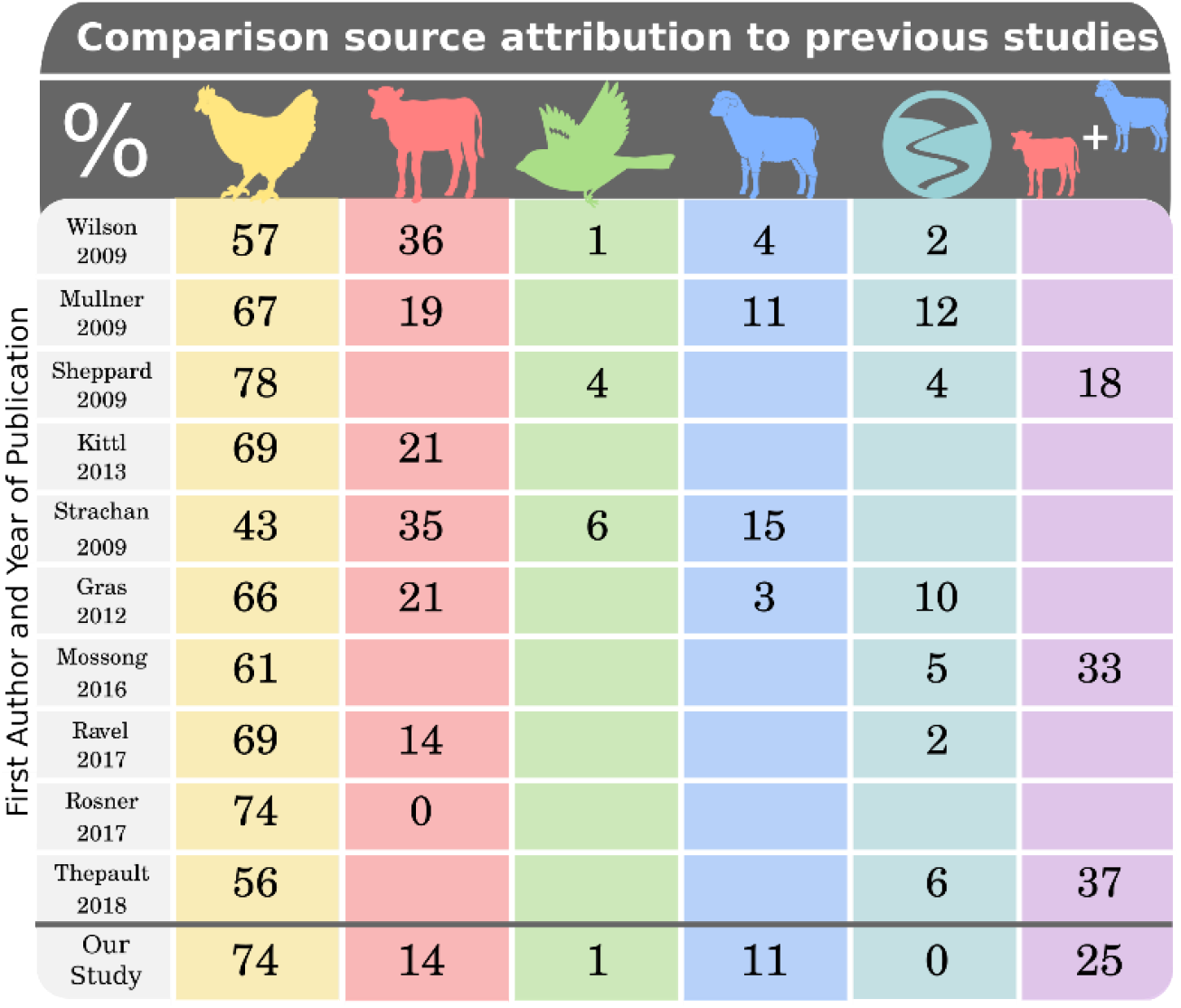
Comparison of our source attribution to previously published studies

### The fine-grained structure of source attribution can be identified with machine learning

Attribution predictions are inferred from the observed frequencies of genotypes in host reservoirs assessed through sampling. However, the relative source composition observed in sampling does not necessarily correspond to host contributions to human infection as some strains that are found at low frequency in the host could be more infectious to humans. For example, some *C. jejuni* strains increase in relative frequency through different stages of the poultry slaughter and production chain because they have genes that promote survival outside of the host (58). There is also evidence that there is a genetic bottleneck at the point of human infection that promotes colonization by strains that have specific genes conferring human niche tropism (59). Analysis of WGS or cgMLST data can potentially allow for changes in relative frequency and provide finer-grained source attribution, potentially at the level of the individual genome.

To identify evidence of differential host affinities, we applied treeBreaker (43) to trace the evolution of a host association along the phylogeny of CC-21, the most commonly found clonal complex in human infection (27). CC-21 frequently colonizes all host sources analysed in this study and is therefore considered a generalist strain, potentially complicating accurate attribution. TreeBreaker detected a change in host association on a branch that groups together a cattle-associated ST-21 subgroup with the cattle-associated lineages ST-982 and ST-806 (Figure 5A). The source composition in this clade (asterisked in Figure 5A) differed from the rest of CC-21, which were predominantly composed of chicken and sheep isolates. Moreover, the asterisked clade differed in its propensity for transmission to humans. Overall, CC-21 was over-represented among human infections, perhaps reflecting its generalist affinities. Yet the asterisked clade was over-represented only 1.7 to 3.6-fold, compared to 5.5 to 6.2-fold for the rest of CC-21 (Figure 5B).

**Figure 5:**
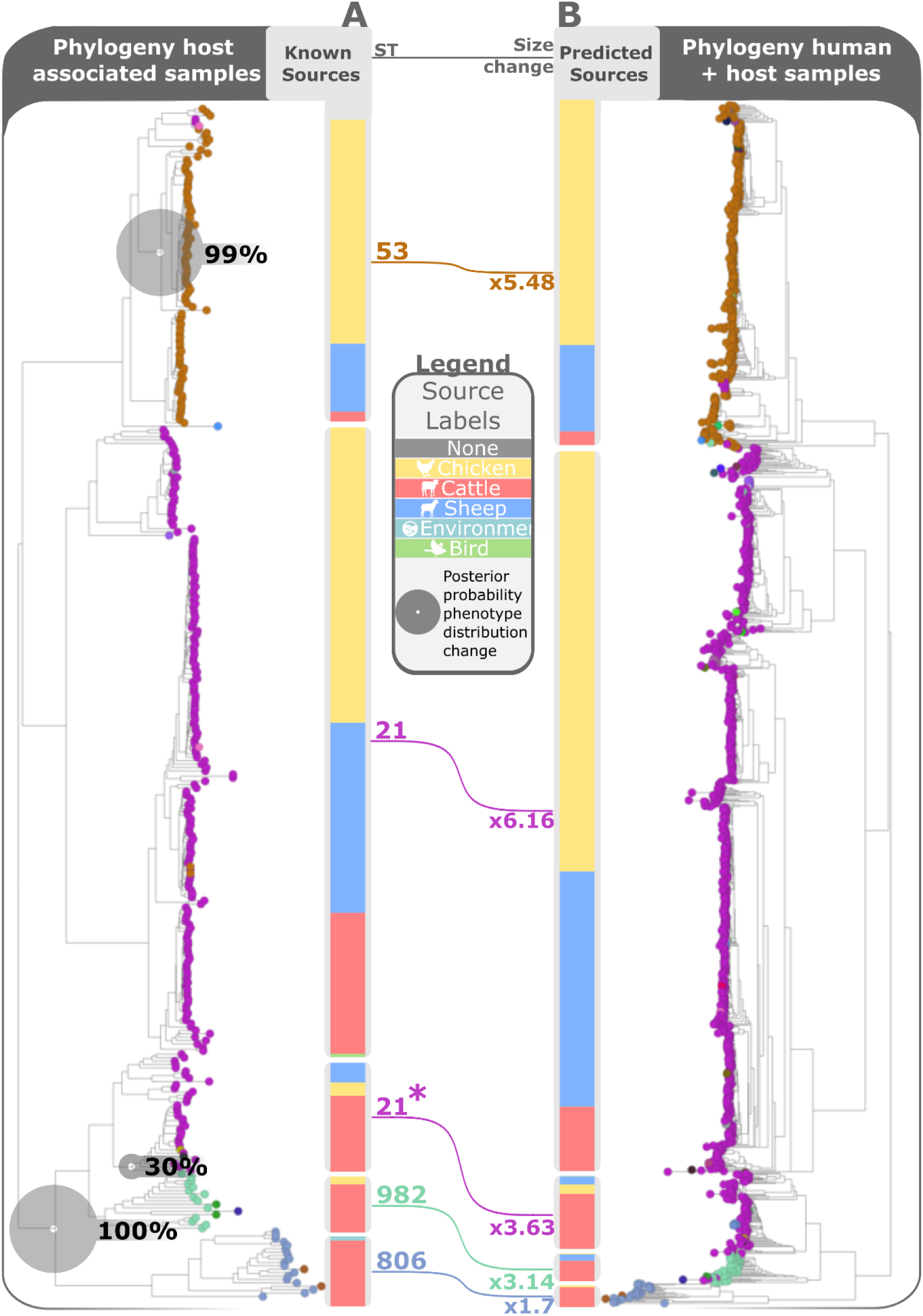
Phylogeny of clonal complex 21 of host animal associated samples (A) and bar charts showing the known source distribution and human samples (B) alongside the predicted source distribution. The phylogeny is based on Neighbour joining using hamming distance of the k-mers drawn from WGS. The connecting lines show the increase in frequency of the clades in human samples and the size of the grey circles show the posterior probability of a change in phenotypic distribution along the branches of the tree.

As the host association changed within CC-21, the ability to transmit to humans appears to have changed as well. This in turn induced a change in the source composition of CC-21 sampled from human infections compared to CC-21 sampled from animals. Previous studies analysing source attribution based on MLST would have overlooked these shifts.

## Outlook and conclusions

The increasing availability of large pathogen genome datasets, algorithms and resources for analysing them, has created possibilities for investigating the transmission of zoonotic diseases that are incompletely understood. It is clear from the data presented here that tree-based ensemble methods for machine learning classification using bacterial genomic data provide considerable utility for improving the accuracy host source attribution for human campylobacteriosis. Key to the effectiveness of this approach is leveraging the full gradient of genomic differentiation afforded by WGS or cgMLST analysis. Host associated genetic variation can be observed in both core and accessory genes (60) but using these data presents practical considerations. With more computational resources available, it may be possible to analyse all k-mers present in the WGS samples (here 109,675,176 unique kmers) with multiple algorithms accompanied by cross-validation and bootstrap replication.

Beyond simple attribution to host source, resolving the fine-grained structure of genomic signatures of association has considerable potential to account for differences in the relative frequency of sub-lineages in samples taken from reservoir hosts and human disease. This can provide important clues about the propensity of strains to survive outside of the host for long enough to transmit to humans as well as the capacity to colonize the human gut given the opportunity (58,59). This of course leads to questions about the genomic basis of bacterial adaptation, specifically the extent to which ‘associated’ genetic elements represent adaptations and whether the same genes and alleles enable colonisation of different host animals.

Improving on the approaches described here, better sampling and incremental training of the XGBoost classifier has considerable potential. The classifier’s low computational requirements and high prediction speed make it an excellent tool for analysing large genome datasets. Furthermore, by using phylogeny-aware train/test splitting for measuring performance, prediction remains accurate when new genetic variants are introduced because the algorithm can be incrementally trained with new data. This has considerable potential for developing automated and continuous disease surveillance systems to reduce campylobacteriosis that remains one of the most common food-borne illness in the world.

## Acknowledgments

N. A. is a recipient of a BBSRC scholarship and thus supported by funding from the Biotechnology and Biological Sciences Research Council (BBSRC) (grant number BB/M011224/1). SKS was supported by Wellcome Trust (088786/C/09/Z) and Medical Research Council (MR/M501608/1 and MR/L015080/1) grants. D. J. W. is supported by a Sir Henry Dale Fellowship jointly funded by the Wellcome Trust and the Royal Society (grant number: 101237/Z/13/B) and by the Robertson Foundation. The research was supported by the National Institute for Health Research (NIHR) Oxford Biomedical Research Centre (BRC). The views expressed are those of the author(s) and not necessarily those of the NHS, the NIHR or the Department of Health.

N.A. would like to thank David Eyre, Christophe Fraser and Alexandra Casey for insightful comments.

Computation used the Oxford Biomedical Research Computing (BMRC) facility, a joint development between the Wellcome Centre for Human Genetics and the Big Data Institute supported by Health Data Research UK and the NIHR Oxford Biomedical Research Centre. The views expressed are those of the author(s) and not necessarily those of the NHS, the NIHR or the Department of Health.

## Supporting information

S1 Table. Metadata of all *Campylobacter* isolates used in this study. Contains the accession numbers, year and country of isolation, source label, generalist index, ST, CC, prediction by our classifier, *Campylobacter species* and whether the samples were used in training or testing.

